# A Prelude to Conservation Genomics: First Chromosome-Level Genome Assembly of a Flying Squirrel (Pteromyini: *Pteromys volans*)

**DOI:** 10.1101/2025.03.20.644356

**Authors:** Gerrit Wehrenberg, Angelika Kiebler, Carola Greve, Núria Beltrán-Sanz, Alexander Ben Hamadou, René Meißner, Sven Winter, Stefan Prost

## Abstract

The Siberian flying squirrel (*Pteromys volans*) represents the only European Pteromyini species. Thus, it is biogeographically unique due to its specialised anatomy and biology as a volant rodent. As a result of habitat fragmentation and destruction, Siberian flying squirrels experience severe and ongoing population declines throughout most of their distribution. While considered *Least Concern* throughout their immense Eurasian distribution, this species is red-listed as *Vulnerable* and even *Critically Endangered* in parts of its range. More knowledge about the population structure and overall biology is needed to improve conservation efforts for this umbrella and flagship species of old-growth boreal forests. Here, we present the first chromosome-level genome assembly of any Pteromyini, represented by *P. volans* (Uoulu_pteVol_1.0). The final assembly has a total length of 2.85 Gbp in 19 chromosome-scale scaffolds with only minor differences in the chromosomal structure compared to other Sciuridae. All chromosome-scale scaffolds show indications for telomeres at both ends; the N50 value and busco as well as *k*-mer completeness scores are high with 157.39 Mbp and 97 – 99 %, respectively, indicating chromosome-level quality of the assembly. Based on whole-genome data from 17 rodent species, *P. volans* clusters according to known evolutionary relationships. Additionally, we present a new 16,511 bp long mitogenome unveiling differences from known conspecific mitogenomes. We propose the utility of the new reference genome for further research and development of conservation-applied genetic methods.

## Introduction

The Siberian flying squirrel (*Pteromys volans* (LINNAEUS, 1758); Figure 1a) is the only native representative of its tribe (Pteromyini BRANDT, 1855) in Europe, with its characteristic patagium enabling a highly arboreal lifestyle, gliding between trees and largely avoiding the ground (Thorington et al., 2012; Koprowski et al., 2016). While being nocturnal, *P. volans* is active throughout the boreal seasons and therefore exposed to dramatic environmental and nutritional changes, adapting its activity and behaviour accordingly (Hokkanen et al., 1977; Törmälä et al., 1980; Hanski, 1998; Hanski et al., 2000). After the Eurasian red squirrel (*Sciurus vulgaris* LINNAEUS, 1758), *P. volans* has the second-largest continuous distribution range of any Sciuromorpha BRANDT, 1855 ranging from Northeast Europe throughout the Eurasian boreal forests until the East Asian Pacific coasts including Manchuria, the Korean Peninsula and the islands of Sakhalin and Hokkaidō southwards limited by arid regions of Central Asia (Thorington et al., 2012; Koprowski et al., 2016). The highest phylogenetic and (mitochondrial as well as nuclear) genetic variability can be found among the East Asian populations (Oshida et al., 2005; Lee et al., 2008; Ito Dos Santos et al., 2024), reflected in comprising all four described and currently accepted subspecies (Thorington et al., 2012; Holden-Musser et al., 2016; Hopkins, 2016; Koprowski et al., 2016). In large portions of its Eurasian range, including the last remaining European populations, it is represented by the nominate subspecies *P. v. volans* (Lee et al., 2008; Thorington et al., 2012; Koprowski et al., 2016; Ito Dos Santos et al., 2024).

**Figure 1.**
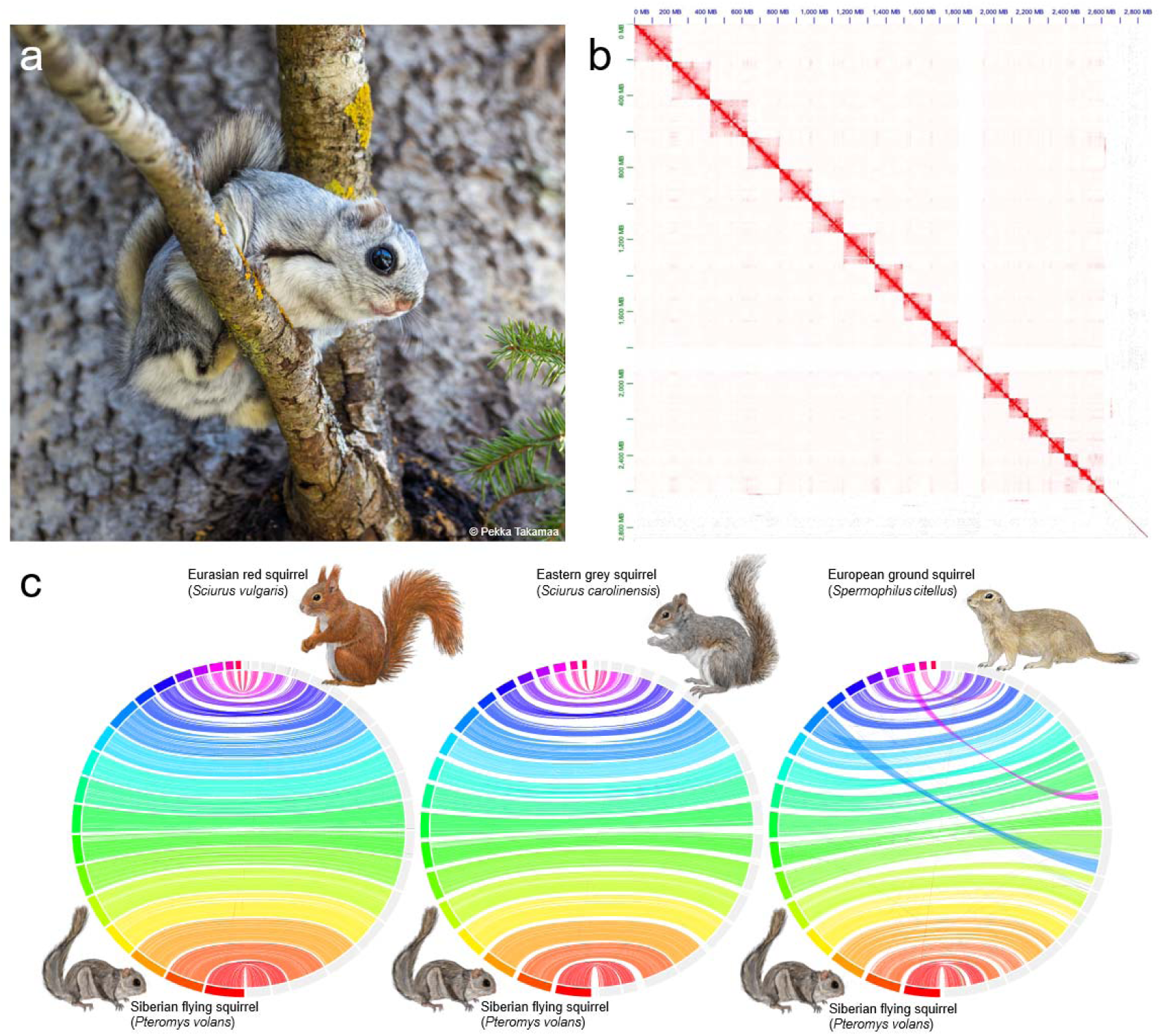
(a) Adult specimen of *Pteromys volans volans* in Lapua, Finland. **(b)** Omni-C contact density map depicting the 19 distinct chromosome-level scaffolds. **(c)** Circos plots comparing the synteny of the chromosome-scale genome assembly of *Pteromys volans* (Uoulu_pteVol_1.0) with chromosome-scale assemblies of a Eurasian red squirrel (*Sciurus vulgaris*), an eastern grey squirrel (*Sciurus carolinensis*), and a European ground squirrel (*Spermophilus citellus*). All three synteny plots are available in a higher resolution in the supplementary information (Supplementary Figures S1-S3).

While its conservation status is globally considered as *Least Concern* by the International Union for Conservation of Nature and Natural Resources (IUCN), populations experience declining trends throughout most of their distribution (Hokkanen et al., 1982; Shar et al., 2016; Zagorodniuk, 2022). Lately, this species has been listed as *Near Threatened* throughout its European distribution (Amori, 2024). Nationally, they are red-listed as *Vulnerable* in Finland (Liukko et al., 2019) and as *Critically Endangered* in Estonia (EELIS, 2025). In fact, the species went extinct in several European countries over the last centuries. Recently, it disappeared in Latvia and Lithuania (Pilāts, 2010, 2023), while a minor population was rediscovered in Northeast Belarus (Abramchuk, 2021). They are legally protected within the European Union (EU) through the Habitats Directive (*Council Directive 92/43/EEC on the Conservation of natural habitats and of wild fauna and flora*) under *Annexes II* and *IV* (European Commission, 2021). On the contrary, it is not continuously protected throughout most of its distribution in Russia (Kurhinen et al., 2011), which represents the majority of its range. Given its relatively large population size, Finland holds special conservation responsibilities within the EU. Regardless of its protection status, its populations are still decreasing mostly due to habitat loss and fragmentation through extensive forestry (Lampila et al., 2009). Additionally, recent changes in Finnish conservation and forestry legislations made the execution of protection of this native rodent ineffective (Santangeli et al., 2013; Wistbacka et al., 2018). Even though Northeast Europe is much less populated by humans compared to the rest of the subcontinent, it is nevertheless highly exploited and thus causes the loss of old-growth mixed forests. The foliage of conifers and cavities in large tree trunks provide cover all year round, while the flying squirrel largely feeds on leaves and catkins of birch and aspen during the warmer season (Hanski et al., 2000; Thorington et al., 2012; Koprowski et al., 2016; Selonen and Wistbacka, 2016). Such dependency on old-growth forests (Mönkkönen et al., 1997; Hanski, 1998) and its charismatic, ecological and phylogenetic uniqueness in Europe makes *P. volans* also an umbrella and flagship species for habitat conservation of intact boreal forests (Hurme et al., 2008; Selonen and Mäkeläinen, 2017). Despite the current conservation focus, the ecology and intraspecific phylogeography of the Siberian flying squirrel is widely unknown.

High-contiguity genome assemblies serve as invaluable references for uncovering such unknown aspects of a species’ biology and advancing applied conservation genetics (Brandies et al., 2019; Paez et al., 2022). Those assemblies not only enhance our understanding of population structures but also enable the identification of informative genetic markers, such as single-nucleotide polymorphisms (SNPs) and microsatellites. With their high contiguity, these reference genomes lay the foundation for developing genetic tools with broad applications in conservation efforts as needed for the Siberian flying squirrel.

While a draft genome assembly of *Glaucomys volans* (LINNAEUS, 1758), native to eastern and southern North America (Koprowski et al., 2016), was published (Wolf et al., 2022), our study provides the first chromosome-level assembly of any Pteromyini. Only such high-contiguity assemblies enable comprehensive analysis of chromosomal structures, including improved SNP localisation, which facilitates linkage disequilibrium mapping and the study of functional diversity in genes, as well as the identification of runs of homozygosity and repeat regions. Here, we present the first high-contiguity, chromosome-level *de novo* genome assembly of *Pteromys volans*.

## Material and Methods

### Sample, DNA extraction and sequencing library preparations

High molecular weight DNA was extracted from muscle tissue taken from a whole-frozen specimen. The specimen died of natural causes and was collected post-mortem at 62.917950°N, 27.631830°E (WGS84) near Kuopio, Finland at the 26^th^ March 2020. It was morphologically assigned as a male of the nominate subspecies of *Pteromys volans volans*. The specimen is stored at -20°C in the collection of the Zoological Museum of the University of Oulu (ZMUO; Collection Management System Kotka/FinBIF: http://id.zmuo.oulu.fi/OV.36069; GBIF: https://www.gbif.org/occurrence/4977508301 (Finbif, 2024)).

We used the *NEB Monarch® HMW DNA Extraction Kit for Cells & Blood* (NEB, USA) for the DNA extraction following the manufacturers protocol. Next, we measured DNA concentration and length utilising the *Qubit dsDNA BR Assay* kit at a Qubit Fluorometer 2.0 (Thermo Fisher Scientific), gel electrophoresis and the *Agilent High Sensitivity DNA* Kit for the *Agilent 2100 Bioanalyzer* (Agilent Technologies, USA). Two *SMRTbell* libraries were prepared according to the instructions of the *SMRTbell Express Prep Kit* v3.0 and sequenced on a *PacBio Revio* system (PacBio, Menlo Park, USA). Next, we generated a proximity ligation sequencing library from the same individual using the *Dovetail Omni-C* Kit (Cantata Bio, USA) following the manufacturers protocol, which was sequenced by *Novogene* in Munich on a *NovaSeq X* platform (Illumina, Inc., San Diego, CA, USA).

### Raw data processing, assembly and scaffolding

First, we extracted the HiFi reads from the BAM file and converted them to the FASTQ format using *BamTools* v2.5.2 (Barnett et al., 2011) with the *-tag “rq”:“>=0.99”* option and *BEDtools* v2.31.1 (Quinlan and Hall, 2010) with the *bamToFastq* option. We then assembled baseline contig-level genome assemblies using the HiFi reads and (1) the *Hifiasm* v0.19.6-r597 assembler (Cheng et al., 2021) and (2) the *Flye* v2.9.4 assembler (Kolmogorov et al., 2019). Based on the *BUSCO* v5.4.7 (Manni et al., 2021) scores and assembly continuity statistics (calculated using *abyss-fac*, (Simpson et al., 2009)) we selected the *Hifiasm* assembly for scaffolding using the Omni-C data. We ran *BUSCO* v5.4.7 in *euk_genome_met* mode and with the gene predictor *metaeuk* (Levy Karin et al., 2020) and the lineage dataset *mammalia_odb10* (which included 24 genomes of 9,226 buscos).

We used Dovetail Genomics’s library QC pipeline (https://omni-c.readthedocs.io/en/latest/index.html) to assess the quality of the Omni-C library. For the Omni-C scaffolding, we applied the Vertebrate Genome Project (VGP) Hi-C scaffolding pipeline (Rhie et al., 2021). In brief, we first filtered the Omni-C reads using the filter_five_end.pl *perl* script (https://github.com/ignacio3437/HiC_mapping_pipeline/tree/master). Next, the reads were merged and PCR duplicates were removed using *Sambamba* v1.0.1 (Tarasov et al., 2015). We used *YaHS* v1.2 (Zhou et al., 2023) for the Omni-C scaffolding and displayed the contact map using *JuicerTools* v1.22.01 (https://github.com/aidenlab/juicertools). Next, we carried out gapclosing using *TGS-GapCloser* v2.0.0 (Xu et al., 2020). We then assessed the scaffolded assembly quality using *BUSCO* v5.4.7 and abyss-fac. We then performed another round of Omni-C scaffolding using the VGP Hi-C scaffolding pipeline and gapclosing using *TGS-GapCloser* v2.0.0, followed by another assembly quality assessment (as described above). Finally, we visualised the generated Omni-C contact map and manually curated the scaffolding using *Juicebox* v2.17.00 (Dudchenko et al., 2018).

### Quality assessment, Repeat and Gene Annotation

The final assembly was assessed with *QUAST* v5.2.0 (Gurevich et al., 2013), *compleasm* v0.2.6, which was shown to be more accurate than *BUSCO* (Huang and Li, 2023), and *BUSCO* v5.4.7, to be able to compare with previous runs. The latter two each utilising the lineage dataset *mammalia_odb10*. In addition, we also investigated the *k*-mer completeness (%; *k* = 21), the consensus quality value (QV; Phred scale), and the *k*-mer multiplicity using *Meryl* v1.4.1 and *Merqury* v1.3 (Rhie et al., 2020). The *k*-mer multiplicity was also esteimented with *GenomeScope* v2.0 (Ranallo-Benavidez et al., 2020). We further estimated the genome-level and scaffold-level coverage using *Qualimap* v2.3 (Okonechnikov et al., 2016) and assessed potential contamination using *BlobTools* v1.1 (Laetsch et al., 2017; Laetsch and Blaxter, 2017) and visualised in *BlobPlots* based on protein sequences found in the database ‘Swiss-Prot’ (release: 04-2024; The UniProt Consortium, 2019) by *diamond* v2.1.6 (Buchfink et al., 2021). To do so, we mapped the HiFi reads onto the Omni-C scaffolded assembly using *minimap2* v2.24 (Li, 2018, 2021). We further identified mitochondrial contigs in the assembly by blasting our assembled mitochondrial genome (outlined in the next section) against the final genome assembly using *NCBI BLAST* v2.16.0 (Boratyn et al., 2013) and removed them from the assembly. The SRY gene (XM_047537220.1) extracted from an eastern grey squirrel genome assembly (*Sciurus carolinensis* GMELIN, 1788; GCA_902686445.2 (Mead et al., 2020a)) was aligned to the novel genome assembly using blastn (*NCBI BLAST* v2.16.0) to determine its genomic location.

To annotate repeats in the genome assembly, we first used *RepeatModeler* v2.0.5 (Flynn et al., 2020) with the search engine *rmblast* v2.14.1+ and *RepeatMasker* v4.1.6 (Smit et al., 2023) with the *Repbase* ‘rodentia’ database (Jurka et al., 2005) and the additional options of: *-nolow* to not hard-mask low complexity DNA or simple repeats and *-gccalc* to calculate the GC content of each contig/scaffold individually. Next, we applied *RepeatMasker* v4.1.6 to the masked genome but with the options: *-xsmall* to change the masking mode from hard to soft-masked, *-noint* to only hard-masks low complex/simple repeats, excluding intronic repeats, and -*gccalc*.

Genes in the masked assembly were predicted based on homology with Gene Model Mapper *GeMoMa* v.1.9 (Keilwagen et al., 2016, 2018) using the following publicly available assemblies at NCBI GenBank: human (*Homo sapiens* LINNAEUS, 1758; GCA_000001405.29), house mouse (*Mus musculus* LINNAEUS, 1758; GCA_000001635.9), eastern grey squirrel (*Sciurus carolinensis*; GCA_902686445.2 (Mead et al., 2020a)), thirteen-lined ground squirrel (*Ictidomys tridecemlineatus* (MITCHILL, 1821); GCA_016881025.1 (Fu et al., 2021)), Arctic ground squirrel (*Urocitellus parryii* RICHARDSON, 1825; GCA_003426925.1), Alpine marmot (*Marmota marmota marmota* (LINNAEUS, 1758); GCA_001458135.2), and groundhog or woodchuck (*Marmota monax* (LINNAEUS, 1758); GCA_021218885.2 (Clarke and Bader, 2024)). The busco completeness of the transcript annotation was assessed with *BUSCO* v5.4.7 utilising the lineage dataset *mammalia_odb10*. Functional annotation of the predicted proteins by *GeMoMa* was conducted by using *diamond* (*blastp* mode) v2.1.6 (Buchfink et al., 2015) with an e-value significance cutoff of ≤ 10^−6^ against the ‘Swiss-Prot’ database (release: 04-2024; The UniProt Consortium, 2019). Furthermore, we annotated gene ontology (GO) terms, domains, and motifs using *InterProScan* v5.73-104.0 (Quevillon et al., 2005; Jones et al., 2014). The gene annotation results were combined utilising modified functional annotation scripts from *MAKER* v3.01.03 (Cantarel et al., 2008; https://github.com/schellt/maker-functional). Gene annotation files are available on Dryad (https://doi.org/10.5061/dryad.3xsj3txth).

Telomere regions were identified using *seqtk* with the option *telo* (https://github.com/lh3/seqtk), and *quarTeT* v1.2.5 (Lin et al., 2023) with the options *-m 10* to *100*.

Lastly, we used the *circlize R* package v0.4.16 (Gu et al., 2014) to perform a six-track circos plot for the largest 19 scaffolds. As an input, we used (A) the telomeres length generated using -m 10, (B) the gene content clustered in 500 kbp windows, (C; D) the repetitive elements (simple and complex repeats) were grouped according to their type (the same TE type found sequentially) and plotted according to their total relative length, and (E) GC content (%; compare Figure 2a).

**Figure 2.**
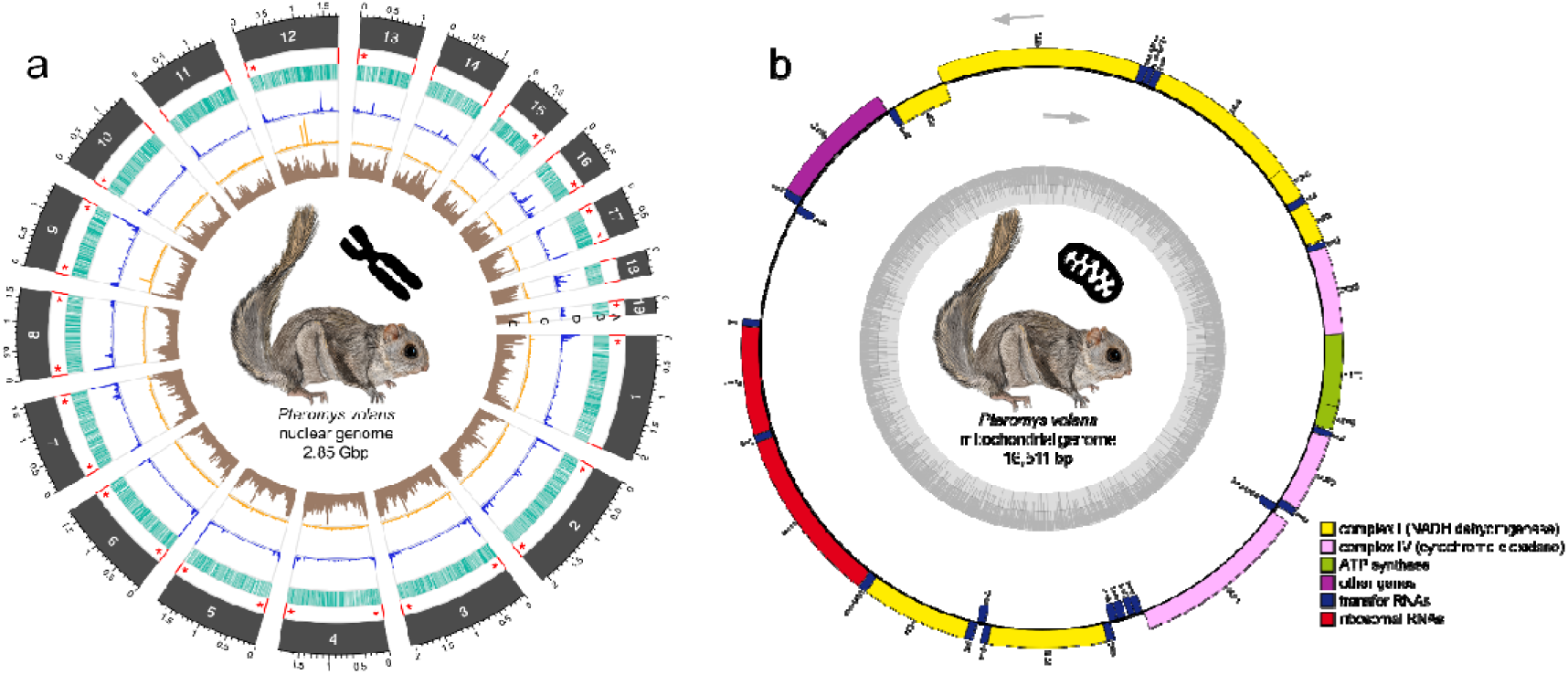
(a) Circos plot of the chromosome-level genome assembly of *Pteromys volans* (Uoulu_pteVol_1.0). The tracks show the following information for 19 scaffolds (length in Gbp) : Red [A]: Position of the telomeres; red asterisks indicate telomeric regions with length of > 100 copies measured with *quarTeT*. Turquoise [B]: Gene content within a 500,000 base pair windows. Blue [C]: Simple repeats. The length of the bars indicates the relative length of the simple repeats after clustering. Orange [D]: Transposable elements (TE) clustered by type. The length of the bars indicates the relative length of the TE after clustering. Brown [E]: GC content. **(b)** Circular gene map of the mitogenome of *Pteromys volans*. The different functional genes groups are shown in different colours, which are shown on the bottom right. The genes transcribed in clockwise and counterclockwise (indicated by arrows) are shown inside and outside of the external circle, respectively. The dark grey colour of inner circle shows the GC content. Figures 2a and 2b as well as the telomere overview plot from *quarTeT* can be found in the Supplementary Information (Supplementary Figures S4 – S6).

### Mitochondrial genome assembly

We used *MitoHifi* v3.2.2 (Uliano-Silva et al., 2023) to reconstruct and the *Flye* v2.9.4 polisher (Kolmogorov et al., 2019) to polish the mitochondrial genome.

Protein-coding genes, rRNAs and tRNAs were annotated and visualised with *Chlorobox* online software tools (https://chlorobox.mpimp-golm.mpg.de/), *GeSeq* (Tillich et al., 2017), and *OGDraw* (Greiner et al., 2019). For *GeSeq*, a circular mitochondrial sequence source was selected, keeping only the best annotation. Annotation via “*BLAT search*” (standard settings) with NCBI GenBank reference MW722951.1 (*P. volans*) and third-party references from five Sciuridae FISCHER DE WALDHEIM, 1817 individuals: southern flying squirrel or assapan (*Glaucomys volans*; NC_050026.1); particolored flying squirrel (*Hylopetes alboniger* (HODGSON, 1836); NC_031847.1), red and white giant flying squirrel (*Petaurista alborufus* (MILNE-EDWARDS, 1870); NC_023922.1), *P. volans* (NC_019612.1 (Ryu et al., 2013)), and Eurasian red squirrel (*Sciurus vulgaris*; NC_002369.1 (Reyes et al., 2000)). GenBank annotation was manually double-checked, and redundant replicates were removed (keeping the version with best alignment scores). For *OGDraw*, a mitochondrial sequence source was selected to visualise the generated GenBank file.

### Synteny and phylogenetic analyses

Next, we assessed genome synteny to the chromosome-level reference genomes for the Eurasian red squirrel (*Sciurus vulgaris*; GenBank: GCA_902686455.2 (Mead et al., 2020b)), the eastern grey squirrel (*Sciurus carolinensis*; GenBank: GCA_902686445.2 (Mead et al., 2020a)), and the European ground squirrel or European souslik (*Spermophilus citellus* LINNAEUS, 1766; GenBank: GCA_964194105.1) using JupiterPlot v1.1 (https://github.com/JustinChu/JupiterPlot) using options: *ng = 101-110*, *m = 10000000*.

We then generated a phylogenetic tree for all available Sciuridae genomes and two Muridae species as an outgroup on the NCBI GenBank: house mouse (*Mus musculus*; GCA_000001635.9), black rat (*Rattus rattus* (LINNAEUS, 1758); GCF_011064425.1 (Rane et al., 2020)), southern flying squirrel or assapan (*Glaucomys volans*; GCA_020662805.1 (Wolf et al., 2022)), Eurasian red squirrel (*Sciurus vulgaris*; GCA_902686455.2 (Mead et al., 2020b), eastern grey squirrel (*Sciurus carolinensis*; GCA_902686445.2 (Mead et al., 2020a), eastern fox squirrel (*Sciurus niger* LINNAEUS, 1758; GCA_020740815.1 (Kang et al., 2021)), unstriped ground squirrel (*Xerus rutilus* (CRETZSCHMAR, 1828); GCA_028644305.1), Cape ground squirrel (*Geosciurus inauris* (ZIMMERMAN, 1780); GCA_004024805.1), Siberian chipmunk (*Eutamias sibiricus* (LAXMANN, 1769); GCA_025594165.1 (Li et al., 2022)), Daurian ground squirrel (*Spermophilus dauricus* BRANDT, 1843; GCA_002406435.1), thirteen-lined ground squirrel (*Ictidomys tridecemlineatus*; GCA_016881025.1 (Fu et al., 2021)), Arctic ground squirrel (*Urocitellus parryii*; GCA_003426925.1), Himalayan marmot (*Marmota himalayana* (HODGSON, 1841); GCA_005280165.1 (Bai et al., 2019)), Alpine marmot (*Marmota marmota marmota*; GCA_001458135.2), Vancouver Island marmot (*Marmota vancouverensis* (SWARTH, 1911); GCA_005458795.1), and yellow-bellied marmot or rock chuck (*Marmota flaviventer* (AUDUBON & BACHMAN, 1841); GCA_003676075.3) using 3,269 busco genes. To do so, we first obtained fasta files for each busco gene using *BUSCO_phylogenomics* (https://github.com/jamiemcg/BUSCO_phylogenomics) with default settings, and we aligned them individually using *Mafft* v7.505 (Katoh and Standley, 2013). Next, we reconstructed phylogenetic trees for each gene separately with *iqtree* v2.3.4 (Minh et al., 2020) utilising a GTR+I+G substitution model. Lastly, we combined the gene trees into a species tree using a multi-species coalescent model using *ASTRAL-III* v5.7.8 (Zhang et al., 2018). Phylogenetic discordance was investigated and visualised using *DiscoVista* (Sayyari et al., 2018).

## Results & Discussion

The 2.8 Gbp large genome of *Pteromys volans* is organised in the 19 largest scaffolds (Figure 1b), representing the expected gonosome and 18 autosomes (2*n* = 38; Oshida and Yoshida, 1996; Oshida et al., 2000; Table 1). Mapping the PacBio HiFi reads back to the genome assembly indicates a mean coverage of 22.6×. Scaffold CM117769.1 shows roughly half the mean genome coverage (12.1×; Supplementary Figure S10, Supplementary Table S3) and minimal interchromosomal contacts (Figure 1b), indicating weak or no interaction with other chromosomes. This along with the chromosome assignment by NCBI GenBank and a blastn-based search of the SRY gene against the novel genome assembly identities this scaffold as the heteromorph gonosomes, which confirms the morphological sex determination of the sequenced specimen. The total sequencing depth was not enough to assemble the Y chromosome. Two runs of PacBio generated in total 64.7 Gbp in filtered HiFi reads that were used for contig genome assembly (run 1: 63.9 Gbp, 7,143,058 reads; run 2: 836.4 Mbp, 90,539 reads). We were able to close two assembly gaps with *TGS-GapCloser*. *Compleasm* scores (99.9% complete, 0.78% duplicated, 0% fragmented and only 0.03% missing genes; Table 2) and *BUSCO* scores (97% complete, 2% duplication, 0.6% fragmented and only 2.4% missing genes; Table 2), a *k*-mer completeness of 97.04% (Consensus QV = 42.7294, corresponding to an error rate of 5.33 × 10⁻C), the *k*-mer multiplicity spectrum (Supplementary Figure S9), and assembly statistics (scaffold-level N50 = 157 Mbp, contig-level N50 = 60Mbp, scaffold-level L90 = 18, contig-level L90 = 73, etc.; Table 1) indicate a high contiguity. The *k*-mer multiplicity corresponds closely to the expected genome coverage of ∼22× (Supplementary Figure S9). *GenomeScope* showed a *k*-mer spectrum of an individual with extremely low heterozygosity, estimated as 0.2%, consistent with a previous study reporting very low heterozygosities for the Finnish population of *P. volans* from which the sequenced individual originates (Ito Dos Santos et al., 2024). The largest 19 scaffolds cover 92% of the assembly and showed at least 10 copies of the telomere repeat TTAGGG on either end. More than 100 copies were found on both ends for nine and at one end for seven additional scaffolds. Only three scaffolds did not show telomeres with more than 100 copies of TTAGGG (Figure 2a, Supplementary Figure S6). The 19 largest scaffolds reached *compleasm* scores of 99.89% complete, 0.78% duplicated, 0.07% fragmented and 0.04% missing genes, only one complete gene less than the full assembly, and *BUSCO* scores equal to the whole assembly, indicating a high completeness of the potential chromosomes. The GC-content with 40.77% lies slightly below eutherian levels of 40.9 – 41.8% (Duret and Galtier, 2009). The homology-based gene prediction with *GeMoMa* identified a total of 27,383 genes and 54,238 transcripts with a *BUSCO* completeness of 91.5% (Supplementary Table S2). More than 97.26% (52,752) of those unique protein query sequences were annotated to sequences from the ‘Swiss-Prot’ database with the *Diamond blastp*-option. The functional annotation with *InterProScan* identified 53,845 unique protein sequences, while 73.3% (39,487) could be assigned to at least one Gene Ontology (GO) term. *Blobtools* identified 17 scaffolds belonging to viruses of the phylum Artverviricota (Supplementary Figure S8), which were removed from the assembly. In total, 970,478,186 bp (34.07%; Table 3) were masked as repeat elements.

**Table 1.**
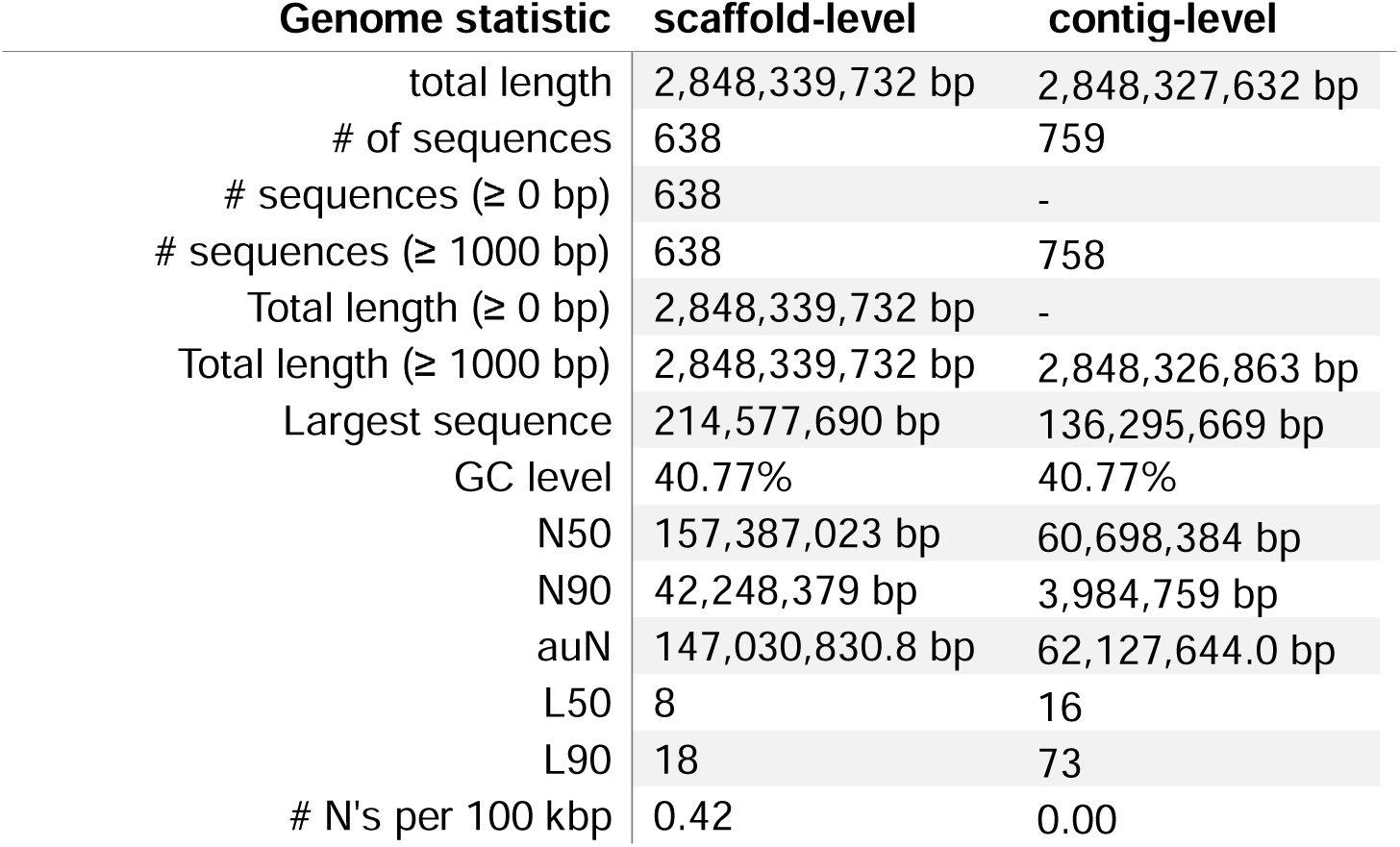
Assembly statistics table on the scaffold-level as well was on the contig-level. (K)bp = (kilo) base pairs; # = number; GC content = Guanine-Cytosine content; auN = area under the Nx curve. A more detailed assembly statistics table can be found in the Supplementary Information (Supplementary Table S1).

**Table 1.**
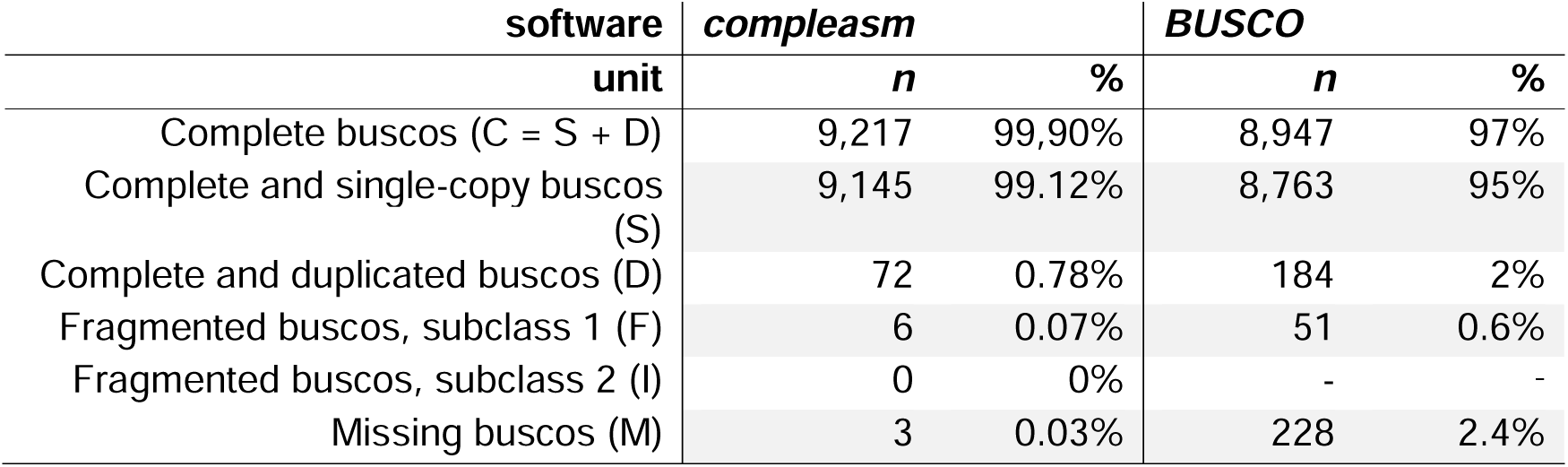
Busco gene scores generated with *compeasm* and *BUSCO* both based on the total number of 9,226 expected orthologs.

| software<br>unit | *compeasm* |  | *BUSCO* |  |
| --- | --- | --- | --- | --- |
|  | *n* | % | *n* | % |
| Complete buscos (C = S + D) | 9,217 | 99.90% | 8,947 | 97% |
| Complete and single-copy buscos (S) | 9,145 | 99.12% | 8,763 | 95% |
| Complete and duplicated buscos (D) | 72 | 0.78% | 184 | 2% |
| Fragmented buscos, subclass 1 (F) | 6 | 0.07% | 51 | 0.6% |
| Fragmented buscos, subclass 2 (I) | 0 | 0% | - | - |
| Missing buscos (M) | 3 | 0.03% | 228 | 2.4% |

**Table 2.** Assembly assignment table estimated by *Repeatmodeler* and *Repeatmasker*. Most repeats fragmented by insertions or deletions have been counted as one element (marked with an asterisk).

| Type of elements | number of elements [n]* | length occupied [bp] | percentage of sequence [%] |
| --- | --- | --- | --- |
| *Retroelements* | 3,255,026 | 886,574,991 bp | 31.13% |
| *SINEs:* | 1,483,628 | 169,453,665 bp | 5.95% |
| *Penelope:* | 62 | 11,402 bp | 0.00% |
| *LINEs:* | 878,225 | 522,064,913 bp | 18.33% |
| *CRE/SLACS* | 0 | 0 bp | 0.00% |
| *L2/CR1/Rex* | 86,215 | 23,004,669 bp | 0.81% |
| *R1/LOA/Jockey* | 0 | 0 bp | 0.00% |
| *R2/R4/NeSL* | 246 | 56,708 bp | 0.00% |
| *RTE/Bov-B* | 5,179 | 1,231,196 bp | 0.04% |
| *L1/CIN4* | 786,498 | 497,749,583 bp | 17.48% |
| *LTR elements:* | 893,173 | 195,056,413 bp | 6.85% |
| *BEL/Pao* | 0 | 0 bp | 0.00% |
| *Ty1/Copia* | 0 | 0 bp | 0.00% |
| *Gypsy/DIRS1* | 6,436 | 1,386,734 bp | 0.05% |
| *Retroviral* | 869,037 | 191,862,047 bp | 6.74% |
| *DNA transposons* | 224,420 | 46,280,608 bp | 1.62% |
| *hobo-Activator* | 163,126 | 32,188,953 bp | 1.13% |
| *Tc1-IS630-Pogo* | 55,186 | 13,103,816 bp | 0.46% |
| *En-Spm* | 0 | 0 bp | 0.00% |
| *MULE-MuDR* | 251 | 41,323 bp | 0.00% |
| *PiggyBac* | 521 | 160,128 bp | 0.01% |
| *Tourist/Harbinger* | 323 | 50,596 bp | 0.00% |
| *Other (Mirage, P-element, Transib)* | 0 | 0 bp | 0.00% |
| *Rolling-circles* | 879 | 195,645 bp | 0.01% |
| *Unclassified:* | 59,407 | 3,502,033 bp | 0.12% |
| *Total interspersed repeats:* |  | 936,369,034 bp | 32.87% |
| *Small RNA:* | 50,111 | 3,274,868 bp | 0.11% |
| *Satellites:* | 49,554 | 2,663,208 bp | 0.09% |
| *Simple repeats:* | 530,473 | 22,897,566 bp | 0.80% |
| *Low complexity:* | 106,166 | 0 bp | 0.19% |

The novel genome assembly phylogenetically clusters with the genomic data of the other published members of Sciuridae as expected based on other studies (Zelditch et al., 2015; Menéndez et al., 2021; Figure 3).

**Figure 3.**
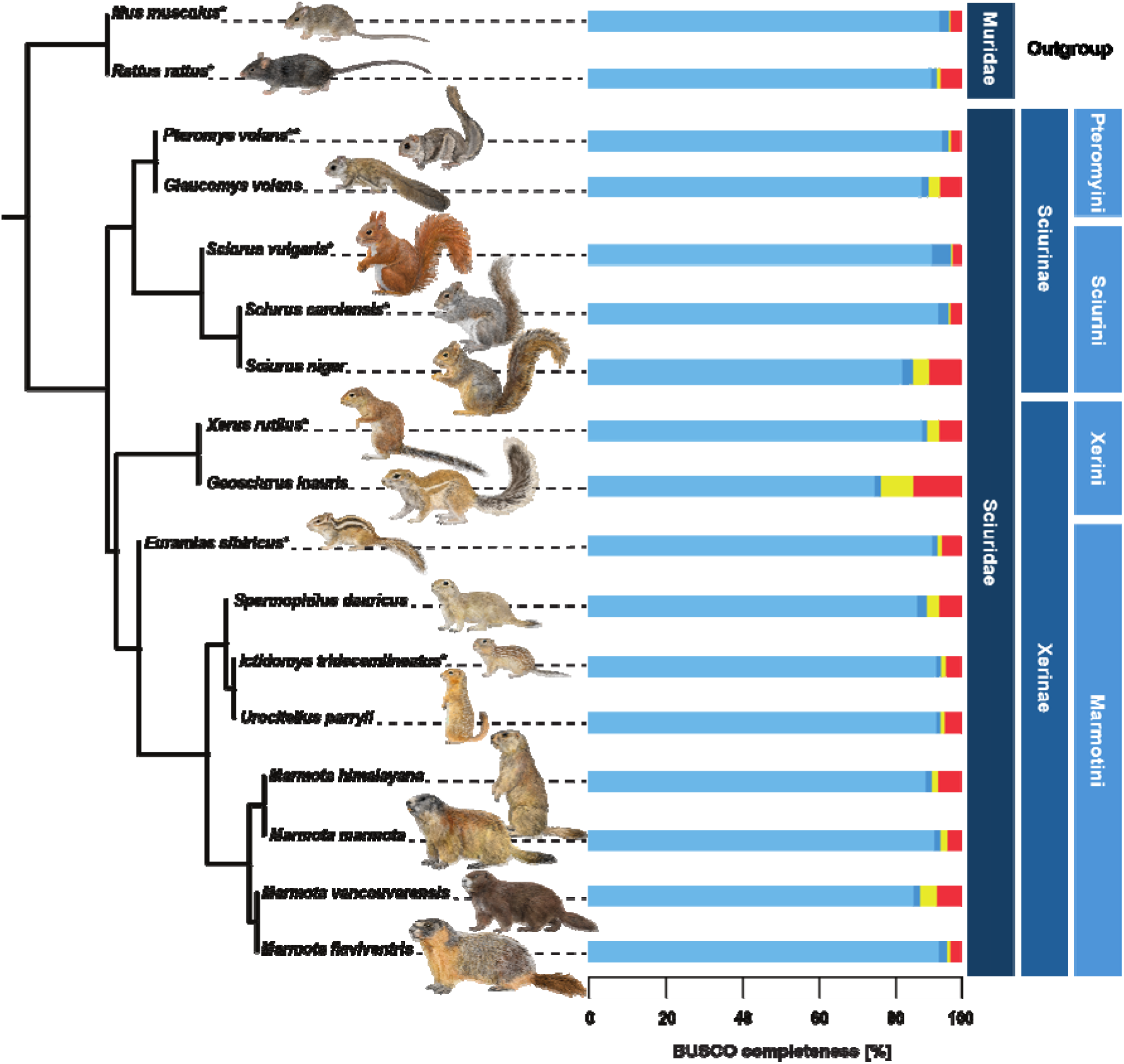
Coalescent phylogenetic tree from 17 rodent species based on 3,269 busco genes each with the assigned busco completeness [%]. Species with available chromosome-level genome assemblies were marked with an asterisk (*). The novel chromosome-level genome assembly from *Pteromys volans* is indicated with two asterisks (**). The outgroup and the according higher taxa are assigned on the right-hand side: families (darkest blue) as well as for the squirrel subfamilies (medium blue) and tribes (lightest blue). The branch quartet frequencies graphs generated by *DiscoVista* can be found in the Supplementary Information (Supplementary Figure S7).

Members of the sister tribe Sciurini LINNAEUS, 1758 showed only minor synteny differences in their chromosomal architecture (Figure 1c). Noteworthy is the apomorph chromosomal fission of scaffold 1 shown in both *Sciurus* species investigated here (Figure 1c). A fused scaffold 1 seems to be a plesiomorph trait kept in *P. volans* if compared to a member of Xerinae OSBORN, 1910 as a clade outside of Sciurinae FISCHER DE WALDHEIM, 1817 (compare to the phylogenetic tree in Figure 3). More distant taxa, such as the European ground squirrel, show a higher degree of chromosomal rearrangements (Figure 1c).

As mentioned above, *Blobtools* identified scaffolds belonging to the viral family Retroviridae. Due to this phylogenetic assignment, those sequences might originate from proviral DNA rather than pararetroviral virions. But we cannot infer if the viruses were active or rather in a proviral phase. Since, the cause for its natural death is unknown and another adult male individual was found with the sequenced specimen, it could shed light on a potential pathogenic relationship of *P. volans* with retroviruses.

The mitochondrial genome has a length of 16,511 bp, including 13 protein-coding genes (CytB, ND1-6, COX1-3, ATP6, and ATP8), two ribosomal RNA genes (12S rRNA and 16S rRNA), and 22 tRNA genes (Figure 2b). This new mitogenome, from the westernmost distribution, contains two fewer base pairs than that of an individual from the Korean Peninsula (Ryu et al., 2013), one of the easternmost ranges, mostly considered to belong to the subspecies *Pteromys volans buechneri* SATUNIN, 1903 (Thorington et al., 2012; Koprowski et al., 2016). Previous genetic studies showed a consistent differentiation between the East Asian mainland populations and the northern Eurasian populations (Oshida et al., 2005; Koh et al., 2008; Ito Dos Santos et al., 2024). However, a more comprehensive sampling and its genomic analysis is required to further resolve the intraspecific phylogeography of the Siberian flying squirrel (Ito Dos Santos et al., 2024), which could be facilitated by the newly presented whole-genome assembly.

For the first time, we provide a chromosome-level genome assembly of a flying squirrel species, namely *Pteromys v. volans* inhabiting extremely fluctuating environments, which will help to understand many aspects of the species’ biology, such as phylogeny, ecological, and nutritional genomics (Ungerer et al., 2008). The sequenced specimen is a representative of the species’ westernmost population, presumably only connected by a narrow corridor to the metapopulation (Kurhinen et al., 2011) with a current conservation focus (Flying Squirrel LIFE; LIFE17 NAT/FI/000469). A previously shown low standing genetic variation within this western population (Ito Dos Santos et al., 2024) is of considerable importance for its adaptive capacity and inferred conservation strategies. Thus, necessary in-depth investigations on inbreeding through runs of homozygosity (ROH) or the genetic health looking at mutational load can only be accomplished with whole-genome assemblies, as provided here. It will also constitute the reference to identify informative genetic markers for conservation applications such as population monitoring. Therefore, this novel genome assembly has the potential to not only shed light on processes of natural selection and adaptation interesting beyond the species context but will provide the basis for applied conservation-relevant studies on the population genetics of the species itself.

## Supporting information

Wehrenberg et al._Supplementary Information

## Data Accessibility Statement

All underlying read data and the assembly are available at NCBI GenBank under BioProject PRJNA1141127. The hard-masked genome assembly and gene annotations are available at Dryad (https://doi.org/10.5061/dryad.3xsj3txth). More detailed results are available in the Supplementary Information.

## Competing Interests Statement

The authors declare no competing interests.

## Author Contributions

**Gerrit Wehrenberg**: Conceptualisation; Data Curation, Formal Analysis; Methodology; Visualisation; Writing – Original Draft Preparation (lead); Writing – Review & Editing (lead). **Angelika Kiebler**: Formal Analysis; Methodology; Visualisation; Writing – Original Draft Preparation; Writing – Review & Editing. **Carola Greve**: Formal Analysis; Methodology; Writing – Review & Editing. **Núria Beltrán-Sanz**: Data Curation, Formal Analysis; Methodology; Visualisation; Writing – Review & Editing. **Alexander Ben Hamadou**: Methodology. **René Meißner**: Formal Analysis; Methodology; Writing – Review & Editing. **Sven Winter**: Conceptualisation; Data Curation, Formal Analysis; Methodology; Supervision; Visualisation; Writing – Review & Editing. **Stefan Prost**: Conceptualisation; Data Curation, Formal Analysis; Funding Acquisition; Methodology; Project Administration; Supervision; Visualisation; Writing – Original Draft Preparation; Writing – Review & Editing.

### Acknowledgments

G.W. and S.P. received funding from Biodiverse Anthropocenes (ANTS) through University of Oulu and the Research Council of Finland PROFI6 funding. We also thank Timo Perätie from Kuopion kaupungin alueelliset ympäristönsuojelupalvelut and the Zoological Museum of the University of Oulu, especially Mikko Vallinmäki, for sampling, sample provision, sample storage and database management. We thank Charlotte Gerheim of the LOEWE Translational Biodiversity Genomics (TBG) project based in the Senckenberg Research Institute and Museum, Frankfurt, for support with lab work. The authors wish to acknowledge CSC – IT Center for Science, Finland, for computational resources. We also thank the Genome Technology Center (RGTC) at Radboud University Medical Center (Radboudumc; Nijmegen, The Netherlands) and the Bioscientia Institut für Medizinische Diagnostik GmbH (Ingelheim, Germany) for providing the PacBio SMRT sequencing service on the PacBio Revio platform. Yannis Schöneberg for helping with the data analysis. We are grateful for the permission to use a wonderful photograph of a Finnish *Pteromys volans volans* by Pekka Takamaa, as well as the scientific illustrations of various rodent species provided by ALADA BOOKS, S.L. Kiitos paljon to Professor Marko Mutanen for the abstract translation into the Finnish language.

## Notes

https://www.ncbi.nlm.nih.gov/bioproject/?term=PRJNA1141127

https://doi.org/10.5061/dryad.3xsj3txth

### Competing Interest Statement

The authors have declared no competing interest.

### Summary of Updates

This revised verion of the manuscript is the latest version after the first peer-review round in Ecology & Evolutuion before resubmitting. There were minor adjustments of the manuscript and additional information on the finctional annotation.

