## Supplementary material for "A Prelude to Conservation Genomics: First Chromosome-Level Genome Assembly of a Flying Squirrel (Pteromyini: *Pteromys volans*)": Wehrenberg et al._Supplementary Information

**Supplementary Figure S4.** Circos plot of the chromosome-level genome assembly of *Pteromys volans* (Uoulu_pteVol_1.0). The tracks show the following information for 19 scaffolds (length in Gbp): Red [A]: Position of the telomeres; red asterisks indicate telomeric regions with length of > 100 copies measured with quarTeT. Turquoise [B]: Gene content within a 500.000 base pair windows. Blue [C]: Simple repeats. The length of the bars indicates the relative length of the simple repeats after clustering. Orange [D]: Transposable elements (TE) clustered by type. The length of the bars indicates the relative length of the TE after clustering. Brown [E]: GC content. The telomere overview plot generated with quarTeT can be found as Supplementary Figure S6. ....................6

**Supplementary Figure S5.** Circular gene map of the mitogenome of *Pteromys volans*. The different functional genes groups are shown in different colours, which are shown on the bottom right. The genes transcribed in clockwise and counterclockwise (indicated by arrows) are shown inside and outside of the external circle, respectively. The dark grey colour of inner circle shows the GC content. ....................7

**Supplementary Figure S6.** Telomere overview plots of the chromosome-level genome assembly of *Pteromys volans* (Uoulu_pteVol_1.0) generated with quarTeT indicating gaps (orange squares) and telomeres (blue triangles). Each plot showcases the results with a different threshold (-m = the min. repeat times to be reported): **(a)** -m10; **(b)** -m20; **(c)** -m40; **(d)** -m60; **(e)** -m80; **(f)** -m100. The most conservative results were presented in the publication (f; indicated by a blue frame).....................8

**Supplementary Figure S7.** Branch quartet frequencies graphs generated by *DiscoVista*. Bars show the relative frequencies of the quartet topologies. The frequency of the species tree topology among gene trees is shown in red, and the other two alternative topologies are shown in blue. The dotted lines indicate the 1/3 threshold. The number of each box indicates the label of the corresponding branch on the tree. On the *x*-axis the exact definition of each quartet topology is shown using the neighbouring branch labels separated by '|'. ....................9

**Supplementary Figure S8.** BlobPlot analysis comparing GC content (*x*-axis). sequencing depth of the PacBio reads (*y*-axis). and taxonomic assignment (phylum) of contigs (colours) show low amounts of viral DNA found in the muscle tissue of *Pteromys volans*. Viral sequences were removed from the final genome assembly. ....................10

**Supplementary Figure S9.** **(a)** Histogram of *k*-mer multiplicity of sequence reads. *k*-mer multiplicity (*x*-axis) is plotted against *k*-mer counts (*y*-axis) to estimate completeness of the novel genome assembly using *Merqury* v1.3. Colours in the plot represent the number of times each *k*-mer is found in the genome assembly. **(b)** Histogram (linear plot) of *k*-mer multiplicity (= coverage; *x*-axis) against *k*-mer counts (= frequency; *y*-axis) generated with *GenomeScope* v2.0. Abbriviations: len = Genome Length [bp]; uniq = unique content [%]; aa = homozygous *k*-mer coverage peak; ab = heterozygous *k*-mer coverage peak; kcov = *k*-mer coverage; err = error rate [%]; dup = duplicated content [%]. Colour code can be found in the legend....................11

**Supplementary Figure S10.** Plot for coverage and GC content across reference assembly generated with *Qualimap* v2.3. Scaffolds are labelled with their assigned Chromosome numbers and divided by dashed lines. The Scaffold 11

### Supplementary Results

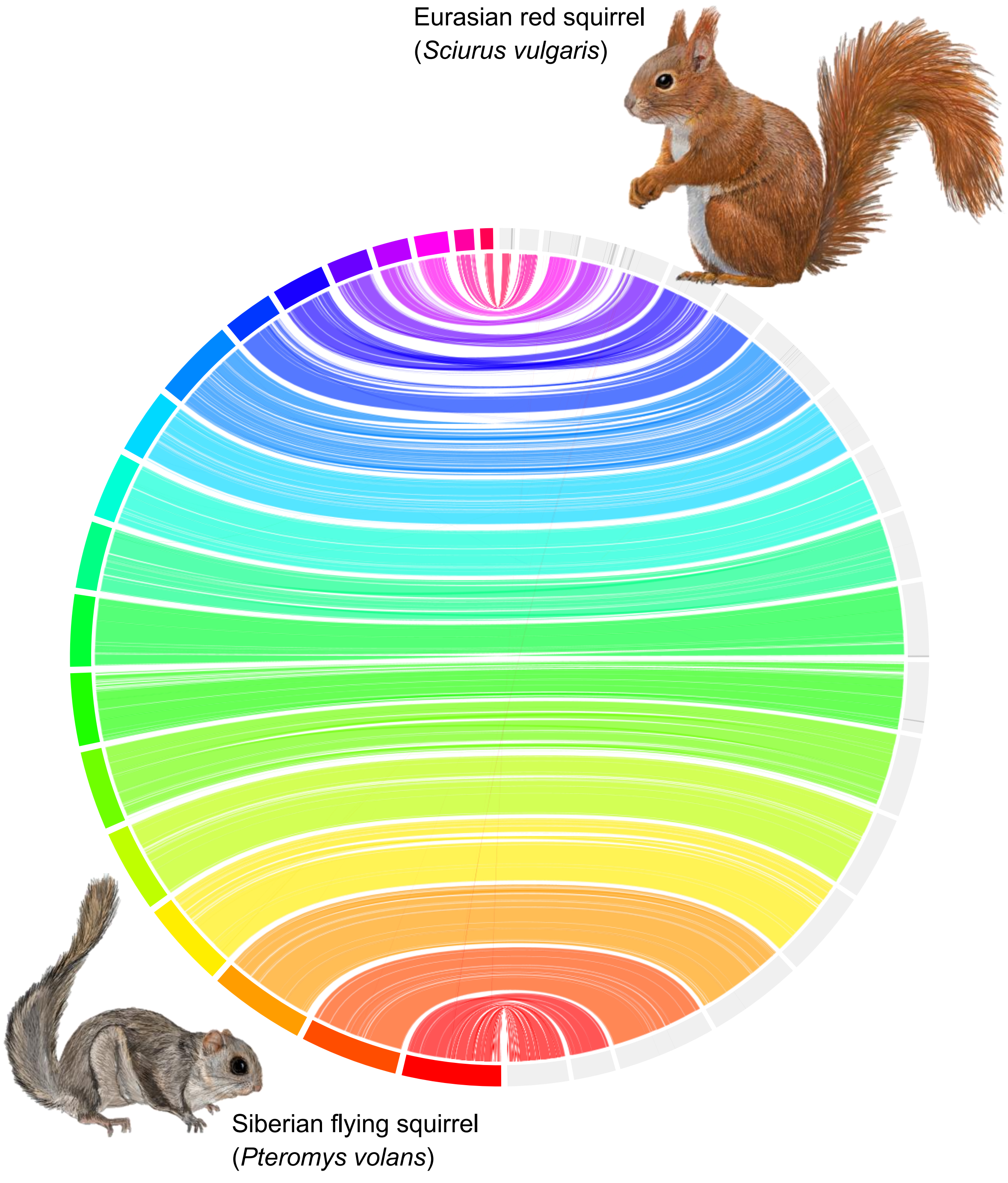

**Supplementary Figure S1.** Circos plot comparing the synteny of the chromosome-scale genome assembly of *Pteromys volans* (Uoulu_pteVol_1.0) with chromosome-scale assemblies of a Eurasian red squirrel (*Sciurus vulgaris* LINNAEUS, 1758).

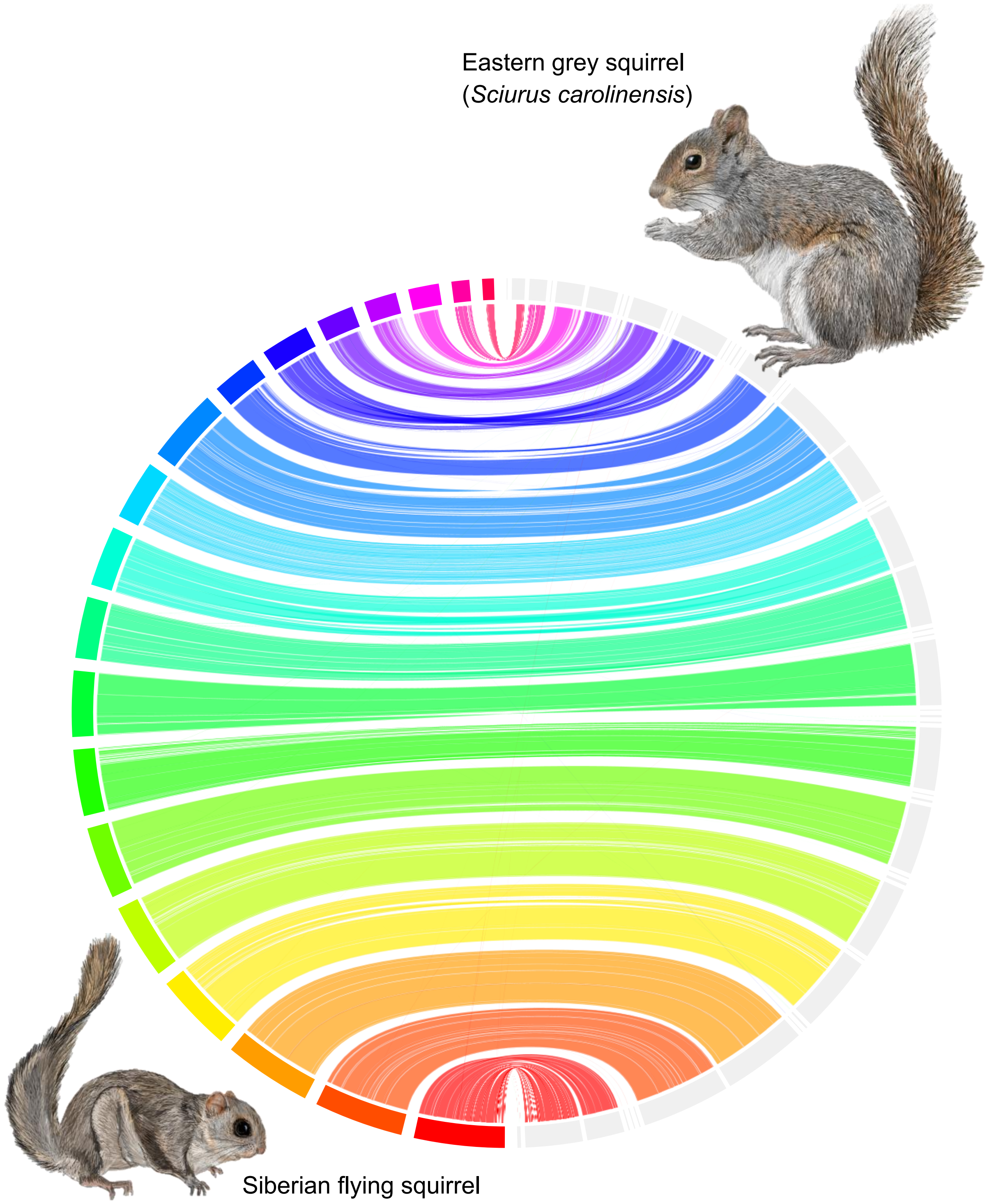

**Supplementary Figure S2.** Circos plots comparing the synteny of the chromosome-scale genome assembly of *Pteromys volans* (Uoulu_pteVol_1.0) with chromosome-scale assemblies of an eastern grey squirrel (*Sciurus carolinensis* GMELIN, 1788).

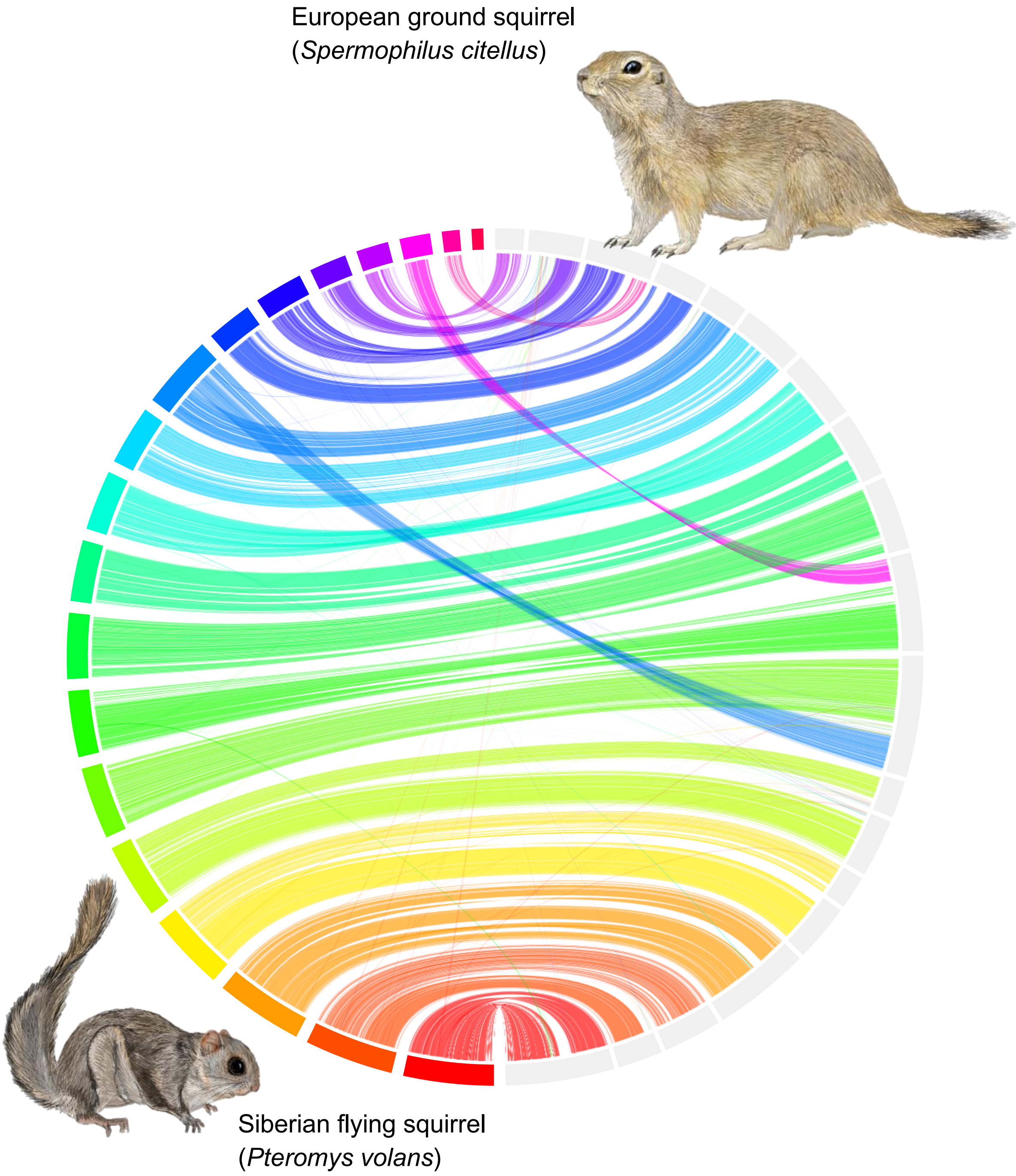

**Supplementary Figure S3.** Circos plots comparing the synteny of the chromosome-scale genome assembly of *Pteromys volans* (Uoulu_pteVol_1.0) with chromosome-scale assemblies of a European ground squirrel (*Spermophilus citellus* (LINNAEUS, 1766)).

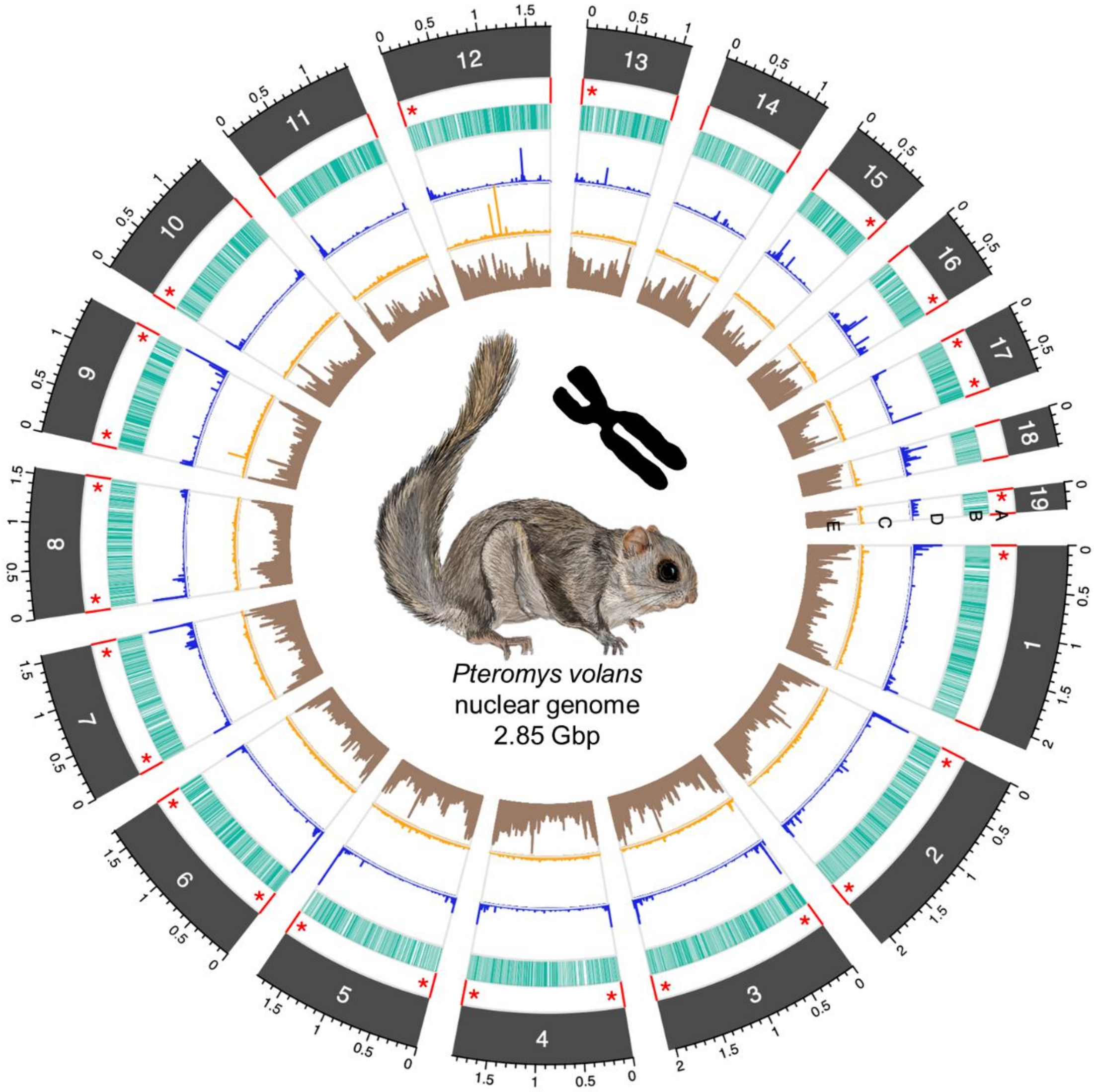

**Supplementary Figure S4.** Circos plot of the chromosome-level genome assembly of *Pteromys volans* (Uoulu_pteVol_1.0). The tracks show the following information for 19 scaffolds (length in Gbp): Red [A]: Position of the telomeres; red asterisks indicate telomeric regions with length of > 100 copies measured with *quarTeT*. Turquoise [B]: Gene content within a 500.000 base pair windows. Blue [C]: Simple repeats. The length of the bars indicates the relative length of the simple repeats after clustering. Orange [D]: Transposable elements (TE) clustered by type. The length of the bars indicates the relative length of the TE after clustering. Brown [E]: GC content. The telomere overview plot generated with *quarTeT* can be found as Supplementary Figure S6.

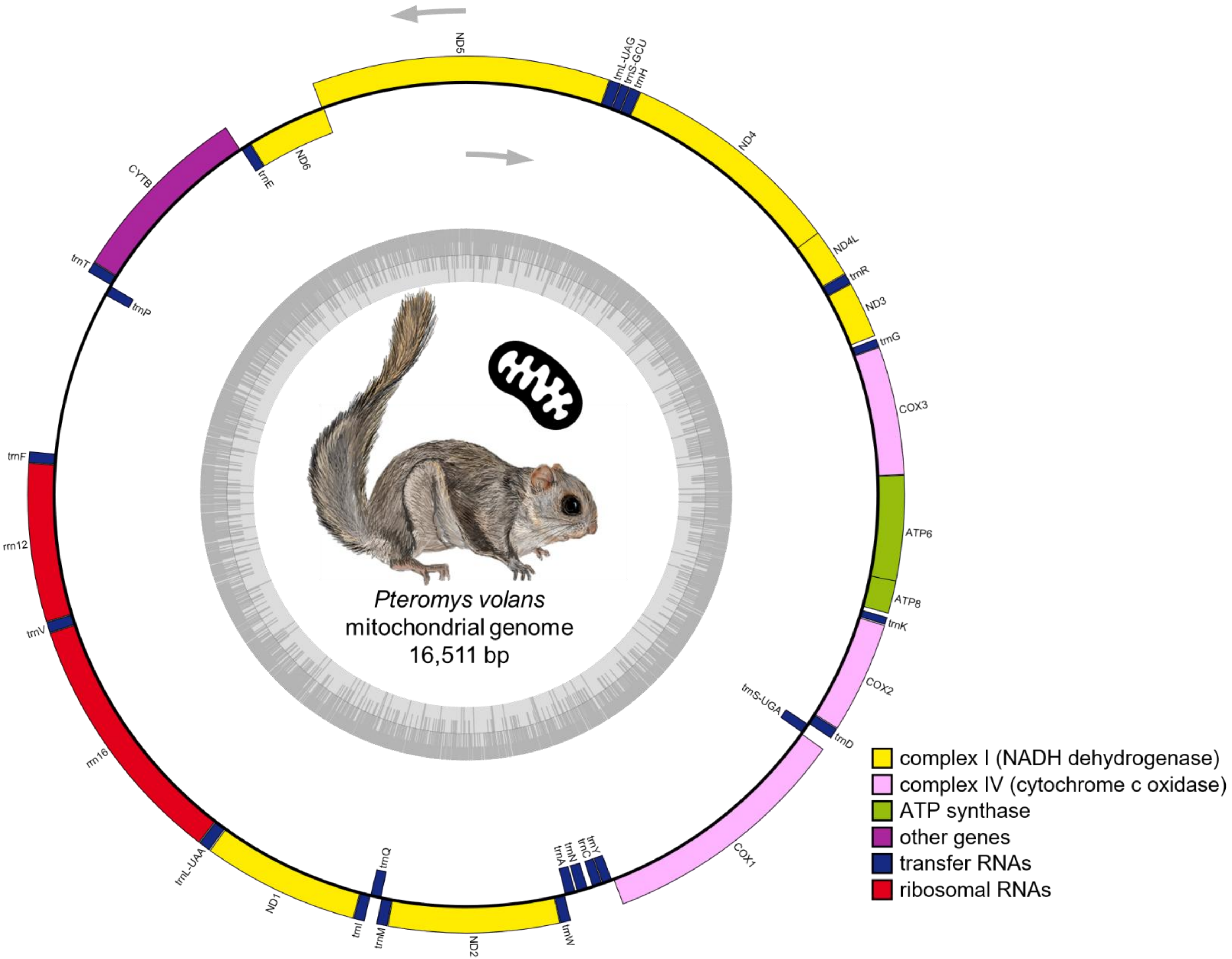

**Supplementary Figure S5.** Circular gene map of the mitogenome of *Pteromys volans*. The different functional genes groups are shown in different colours, which are shown on the bottom right. The genes transcribed in clockwise and counterclockwise (indicated by arrows) are shown inside and outside of the external circle, respectively. The dark grey colour of inner circle shows the GC content.

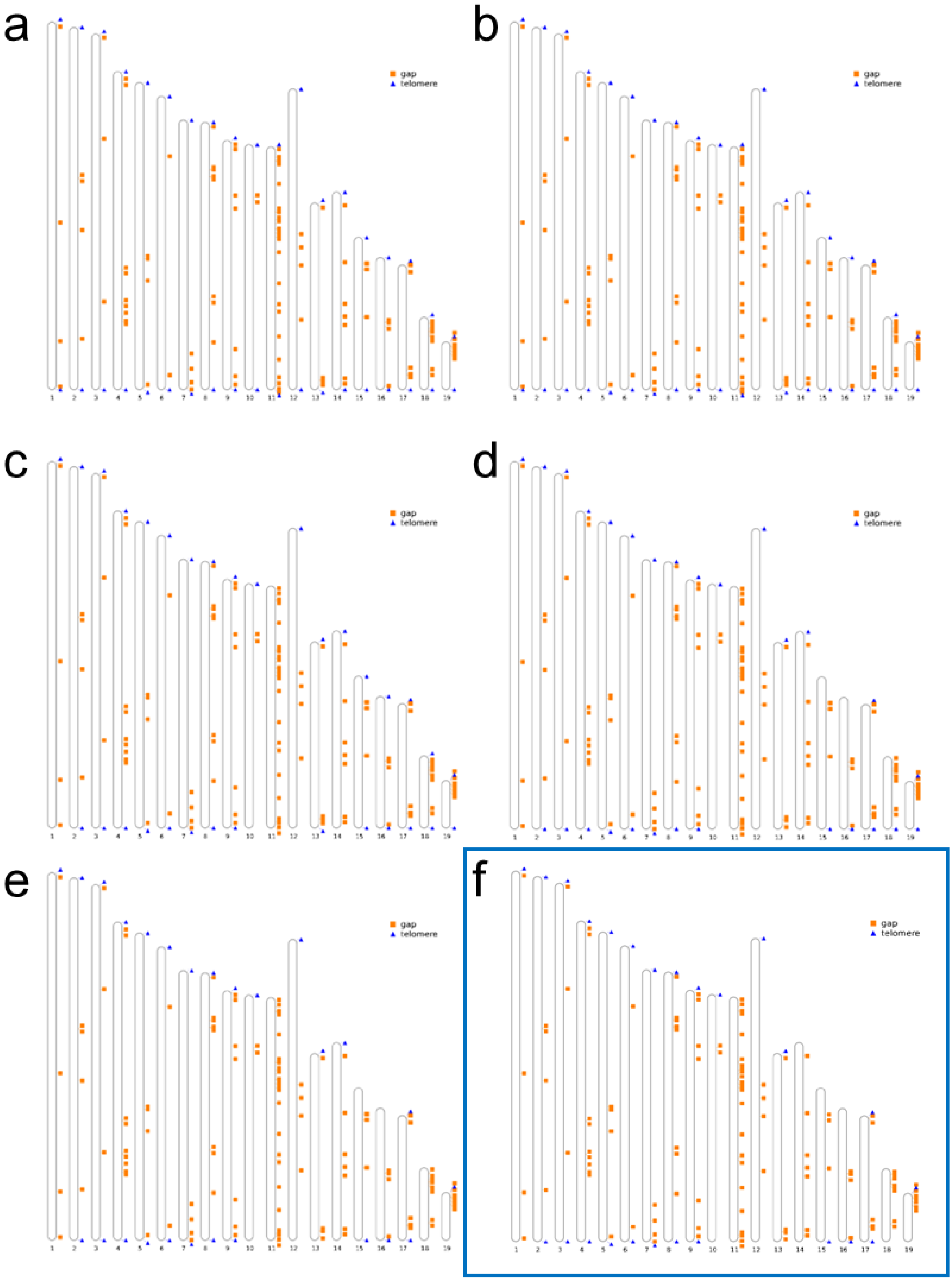

**Supplementary Figure S6.** Telomere overview plots of the chromosome-level genome assembly of *Pteromys volans* (Uoulu_pteVol_1.0) generated with *quarTeT* indicating gaps (orange squares) and telomeres (blue triangles). Each plot showcases the results with a different threshold (*-m* = the min. repeat times to be reported): **(a)** *-m10*; **(b)** *-m20*; **(c)** *-m40*; **(d)** *-m60*; **(e)** *-m80*; **(f)** *-m100*. The most conservative results were presented in the publication (f; indicated by a blue frame).

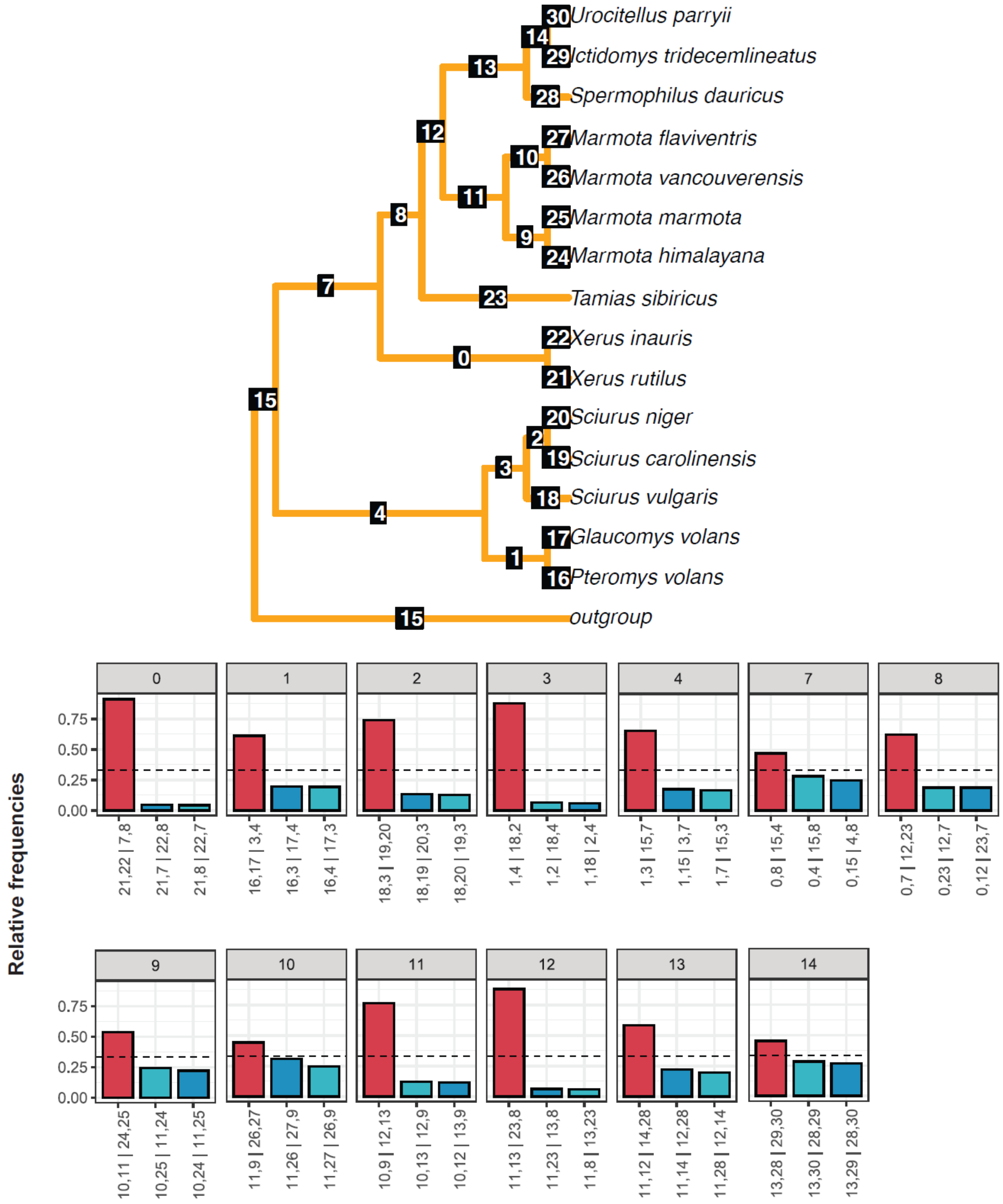

**Supplementary Figure S7.** Branch quartet frequencies graphs generated by *DiscoVista*. Bars show the relative frequencies of the quartet topologies. The frequency of the species tree topology among gene trees is shown in red, and the other two alternative topologies are shown in blue. The dotted lines indicate the 1/3 threshold. The number of each box indicates the label of the corresponding branch on the tree. On the x-axis the exact definition of each quartet topology is shown using the neighbouring branch labels separated by '|'.

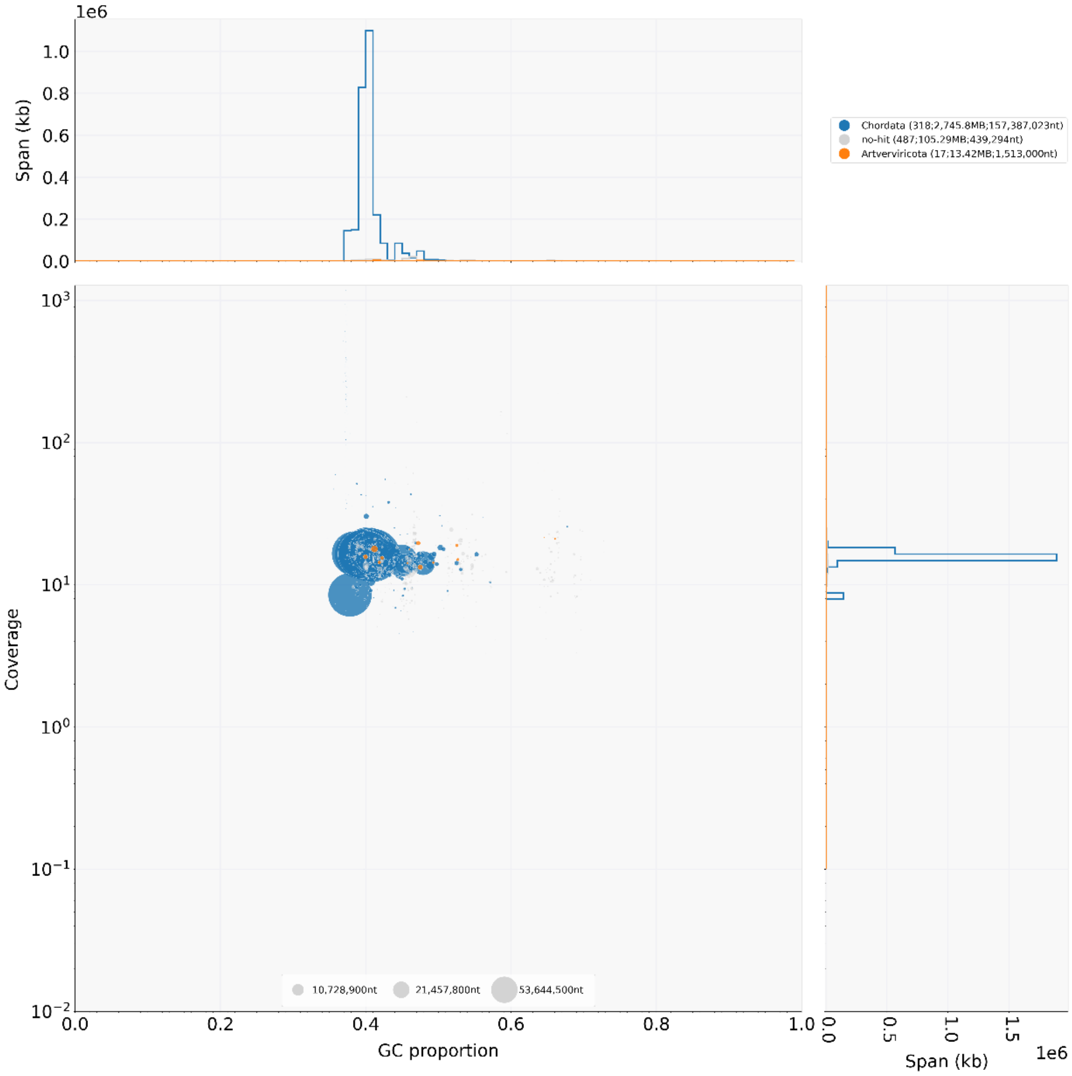

**Supplementary Figure S8.** BlobPlot analysis comparing GC content (*x*-axis). sequencing depth of the PacBio reads (*y*-axis). and taxonomic assignment (phylum) of contigs (colours) show low amounts of viral DNA found in the muscle tissue of *Pteromys volans*. Viral sequences were removed from the final genome assembly.

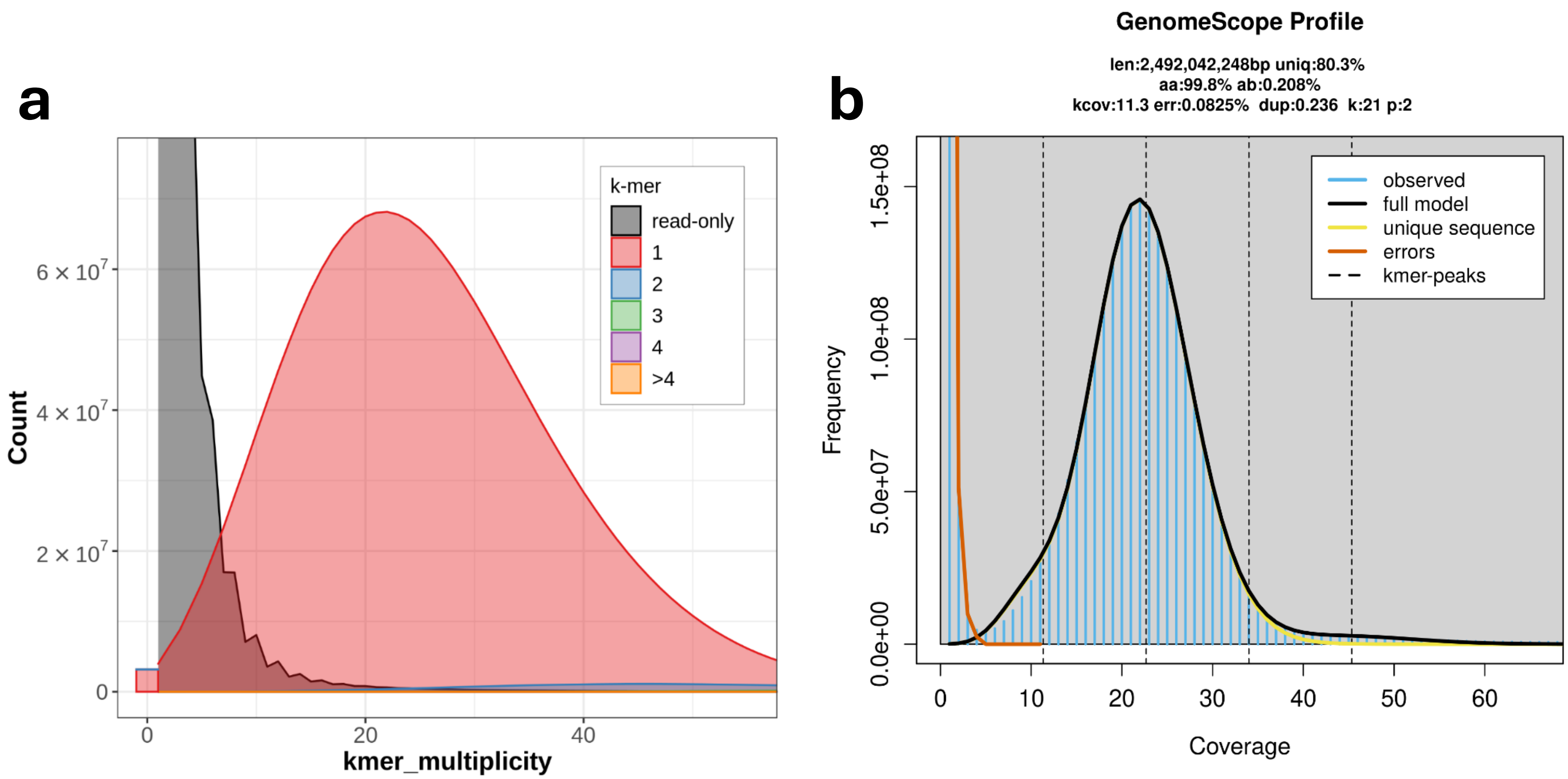

**Supplementary Figure S9. (a)** Histogram of *k*-mer multiplicity of sequence reads. *k*-mer multiplicity (*x*-axis) is plotted against *k*-mer counts (*y*-axis) to estimate completeness of the novel genome assembly using *Merqury* v1.3. Colours in the plot represent the number of times each *k*-mer is found in the genome assembly. **(b)** Histogram (linear plot) of *k*-mer multiplicity (= coverage; *x*-axis) against *k*-mer counts (= frequency; *y*-axis) generated with *GenomeScope* v2.0. Abbriviations: len = Genome Length [bp]; uniq = unique content [%]; aa = homozygous *k*-mer coverage peak; ab = heterozygous *k*-mer coverage peak; kcov = *k*-mer coverage; err = error rate [%]; dup = duplicated content [%]. Colour code can be found in the legend.

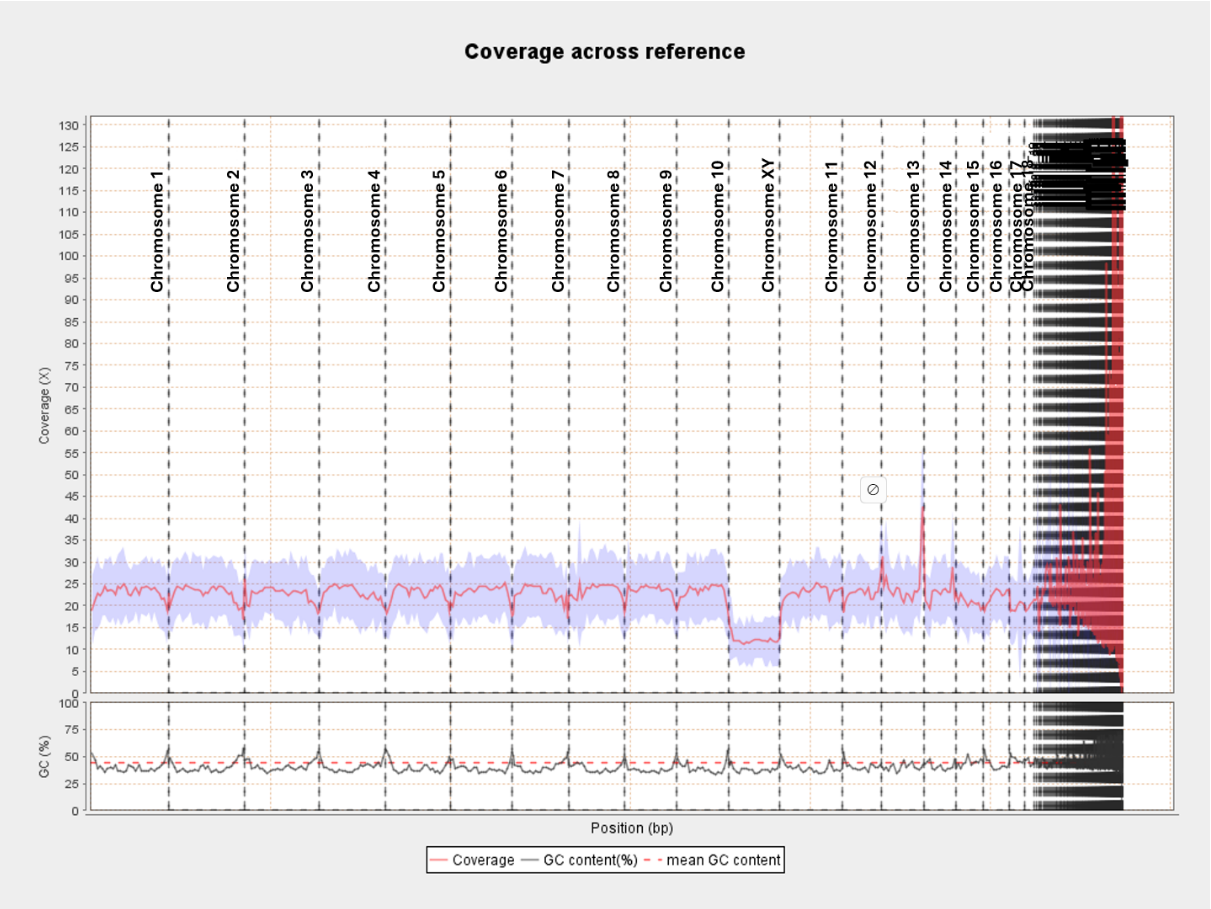

**Supplementary Figure S10.** Plot for coverage and GC content across reference assembly generated with *Qualimap* v2.3. Scaffolds are labelled with their assigned Chromosome numbers and divided by dashed lines. The Scaffold 11 (CM117769.1) shows roughly half the coverage of the other major scaffolds. Along with the results of the contact map (Figure 1b), the chromosome assignment by NCBI and blast results, this indicates that this scaffold is the gonosome.

**Supplementary Table S1.** Assembly statistics table on the scaffold-level as well was on the contig-level. (K)bp = (kilo) base pairs; # = number; GC content = Guanine-Cytosine content; auN = area under the Nx curve.

| Genome statistic | scaffold-level | contig-level |
|---|---|---|
| total length | 2,848,339,732 bp | 2,848,327,632 bp |
| # of sequences | 638 | 759 |
| # sequences (≥ 0 bp) | 638 | - |
| # sequences (≥ 1000 bp) | 638 | 758 |
| # sequences (≥ 5000 bp) | 636 | 749 |
| # sequences (≥ 10000 bp) | 633 | 746 |
| # sequences (≥ 25000 bp) | 582 | 691 |
| # sequences (≥ 50000 bp) | 523 | 623 |
| Total length (≥ 0 bp) | 2,848,339,732 bp | - |
| Total length (≥ 1000 bp) | 2,848,339,732 bp | 2,848,326,863 bp |
| Total length (≥ 5000 bp) | 2,848,337,732 bp | 2,848,312,932 bp |
| Total length (≥ 10000 bp) | 2,848,314,129 bp | 2,848,289,329 bp |
| Total length (≥ 25000 bp) | 2,847,400,174 bp | 2,847,310,491 bp |
| Total length (≥ 50000 bp) | 2,845,303,824 bp | 2,844,845,673 bp |
| Largest sequence | 214,577,690 bp | 136,295,669 bp |
| GC level | 40.77% | 40.77% |
| N50 | 157,387,023 bp | 60,698,384 bp |
| N90 | 42,248,379 bp | 3,984,759 bp |
| auN | 147,030,830.8 bp | 62,127,644.0 bp |
| L50 | 8 | 16 |
| L90 | 18 | 73 |
| # N's per 100 kbp | 0.42 | 0.00 |

**Supplementary Table S2.** Busco gene scores were calculated using annotated transcripts using *BUSCO* (54,238 protein sequences; 27,383 coding DNA sequences (CDS)) based on the total number of 9,226 expected orthologs (lineage data set *mammalia_odb10* including 24 genomes).

| | Protein sequences | | CDS | |
|---|---|---|---|---|
| unit | n | % | n | % |
| Complete BUSCOs (C = S + D) | 8,435 | 91.5% | 8,433 | 91.5% |
| Complete and single-copy BUSCOs (S) | 4,701 | 51.0% | 8,216 | 89.1% |
| Complete and duplicated BUSCOs (D) | 3,734 | 40.5% | 217 | 2.4% |
| Fragmented BUSCOs(F) | 301 | 3.3% | 235 | 2.5% |
| Missing BUSCOs (M) | 490 | 5.2% | 558 | 6.0% |

**Supplementary Table S3.** Per scaffold statistics showing the scaffold length [bp], mapped bases, mean coverage, and the coverage standard deviation generated with *Qualimap* v2.3. Scaffold 11 (CM117769.1; assigned as gonosome) shows a considerably lower mean coverage and coverage standard deviation. Mean coverage for the whole assembly is 22.6166×.

| Chromosome | Scaffold | GenBank ID | Length [bp] | Mapped bases | Mean Coverage | Standard deviation |
|---|---|---|---|---|---|---|
| 1 | 1 | CM117751.1 | 214,577,690 | 4,950,981,021 | 23.07314 | 5.400098 |
| 2 | 2 | CM117752.1 | 211,549,263 | 4,852,606,657 | 22.938424 | 5.672579 |
| 3 | 3 | CM117753.1 | 207,559,492 | 4,709,923,921 | 22.691923 | 5.589127 |
| 4 | 4 | CM117754.1 | 185,676,949 | 4,264,990,996 | 22.969954 | 5.508416 |
| 5 | 5 | CM117755.1 | 179,342,594 | 4,133,582,513 | 23.048526 | 5.488774 |
| 6 | 6 | CM117756.1 | 171,299,418 | 3,995,538,289 | 23.32488 | 5.409834 |
| 7 | 7 | CM117757.1 | 157,387,023 | 3,608,255,475 | 22.926004 | 5.688032 |
| 8 | 8 | CM117758.1 | 156,106,043 | 3,689,451,166 | 23.634262 | 6.34366 |
| 9 | 9 | CM117759.1 | 145,542,709 | 3,425,100,506 | 23.533302 | 5.643468 |
| 10 | 10 | CM117760.1 | 143,035,200 | 3,342,220,240 | 23.366418 | 5.626549 |
| 11 | 12 | CM117761.1 | 175,561,277 | 4,073,826,765 | 23.204586 | 5.658423 |
| 12 | 13 | CM117762.1 | 108,950,874 | 2,494,628,853 | 22.896823 | 5.777009 |
| 13 | 14 | CM117763.1 | 115,363,772 | 2,798,063,227 | 24.254263 | 8.176572 |
| 14 | 15 | CM117764.1 | 88,932,773 | 2,045,808,666 | 23.003991 | 6.468686 |
| 15 | 16 | CM117765.1 | 77,077,429 | 1,612,197,034 | 20.916591 | 5.341857 |
| 16 | 17 | CM117766.1 | 72,818,191 | 1,619,908,847 | 22.245936 | 5.866627 |
| 17 | 18 | CM117767.1 | 42,248,379 | 835,690,026 | 19.780404 | 8.369105 |
| 18 | 19 | CM117768.1 | 27,885,036 | 559,159,382 | 20.05231 | 5.550514 |
| X | 11 | CM117769.1 | 141,649,903 | 1,718,619,800 | 12.13287 | 3.956911 |
